# Transient uterine hypercontractility causes fetal cerebral oxidative stress and enduring mitochondrial and behavioral abnormalities in adolescent male rat offspring

**DOI:** 10.1101/689927

**Authors:** Arvind Palanisamy, Tusar Giri, Jia Jiang, Annie Bice, James D. Quirk, Sara B. Conyers, Susan E. Maloney, Nandini Raghuraman, Adam Q. Bauer, Joel R. Garbow, David F. Wozniak

## Abstract

The impact of transient ischemic-hypoxemic insults on the developing fetal brain is poorly understood despite evidence suggesting an association with neurodevelopmental disorders such as schizophrenia and autism. To address this, we designed an aberrant uterine hypercontractility paradigm with oxytocin to better assess the consequences of acute, but transient, placental ischemia-hypoxemia in term pregnant rats. Using MRI imaging, we confirmed that oxytocin-induced aberrant uterine hypercontractility significantly compromised uteroplacental perfusion. This was supported by the observation of oxidative stress and increased lactate concentration in the fetal brain. Genes related to oxidative stress pathways were significantly upregulated in male, but not female, offspring 1 h after oxytocin-induced placental ischemia-hypoxemia. Persistent upregulation of select mitochondrial electron transport chain complex proteins in the anterior cingulate cortex of adolescent male offspring suggested that this sex-specific effect was enduring. Functionally, offspring exposed to oxytocin-induced uterine hypercontractility showed male-specific abnormalities in social behavior with associated region-specific changes in gene expression and functional cortical connectivity. Our findings, therefore, indicate that even transient but severe placental ischemia-hypoxemia could be detrimental to the developing brain and point to a possible mitochondrial link between intrauterine asphyxia and neurodevelopmental disorders.

## Introduction

Epidemiological studies suggest a link between obstetric complications and neurodevelopmental disorders (NDD),^1–5^ but the underlying biological mechanisms remain largely unexplored. The consequences of severe birth asphyxia are well understood both in human and animal studies,^6–12^ but the neurodevelopmental impact of transient and less severe hypoxemic insults, which are more common during labor and delivery, is poorly understood. For example, aberrant uterine contractility (tachysystole and tetanic uterine contraction) can often lead to transient fetal distress which may or may not necessitate emergent delivery. These episodes are more common, occurring in approximately 5-10% of all laboring women,^13,14^ especially when oxytocin (OXT) is used for induction and augmentation of labor. It is widely presumed that these episodes are usually benign because the surrogate markers of fetal well-being, such as umbilical arterial lactate and pH, are typically unaffected. Though this is reassuring, it is not known if these transient ischemic-hypoxemic episodes have an impact on the fetal brain, arguably the most important organ of interest. This knowledge is critical because of the exquisite sensitivity of the fetal brain to hypoxia, and its susceptibility to oxidative stress because of the lack of robust antioxidant defense mechanisms.^15^ Current animal models that examine this question are contrived because the procedures devised to create placental ischemia-hypoxemia are very invasive (eg., uterine artery ligation or occlusion),^16,17^ require prolonged anesthesia, and rarely reflect the transient and reversible ischemic-hypoxemic insults that are far more common during labor and delivery. Here, we created a pragmatic and translationally relevant pregnant rat model by stimulating uterine hypercontractility with OXT. We show that such uterine hypercontractility causes transient placental ischemia-hypoxemia, significant oxidative stress in the fetal brain, enduring brain-region specific dysfunction of the mitochondrial electron transport chain complex, abnormal cortical functional connectivity, and subtle neurobehavioral abnormalities especially in the adolescent male offspring.

## Results

### OXT-induced uterine hypercontractility causes significant placental ischemia

Though aberrant uterine contractility is known to cause uteroplacental hypoperfusion in pregnant women, it was unclear whether this could be replicated in a pregnant rat model. Therefore, we addressed this by quantifying the effect of OXT-induced uterine hypercontractility on placental perfusion in term pregnant Sprague Dawley rats (E21) with dynamic contrast-enhanced MRI (**Figure 1A**). Placental uptake of Dotarem^®^ contrast was approximately 45% lower after OXT-induced hypercontractility, indicating a significant decrease in uteroplacental blood flow. Next, we assessed whether the impaired uteroplacental perfusion was accompanied by placental hypoxemia with R2* mapping in a separate set of experiments (**Figure 1B**). Thirty min after OXT, there was a significant change in the junctional zone contrast such that the placentas appeared completely bright with a significantly lower R2*. Though hypoxic placenta is expected to demonstrate a higher R2*, this change in junctional zone contrast is the effect of a longer T2 (and/or T2*), which would reasonably be explained by a significant decrease in iron (or deoxyhemoglobin). This suggested that these placentae were either not receiving much blood or were having it expelled out of them by the tetanic uterine contraction. Collectively, our placental imaging data confirm that OXT-induced hypercontractility causes significant impairment of placental perfusion.

**Figure 1.**
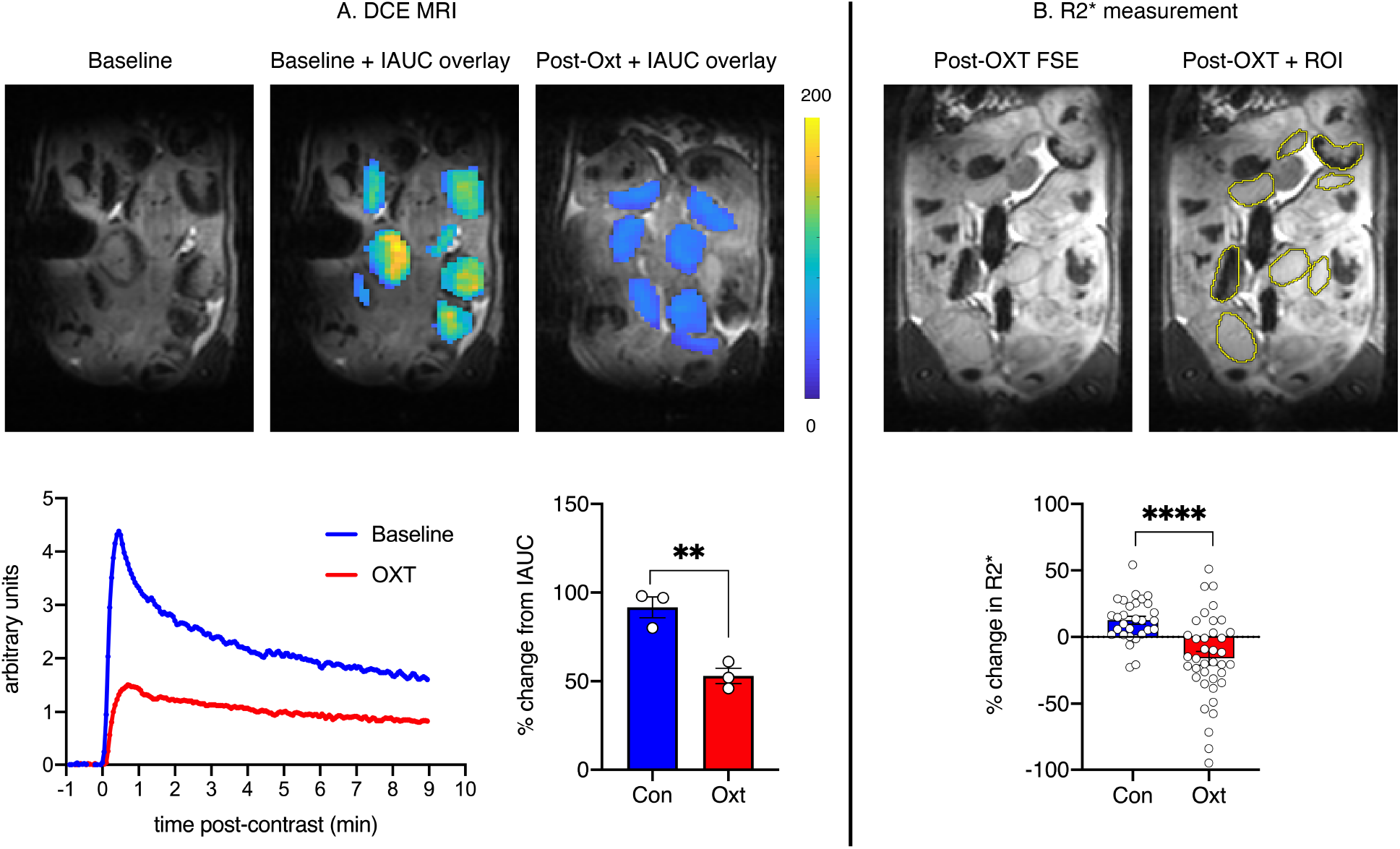
**A) Placental perfusion is significantly impaired after OXT-induced aberrant uterine contractility.** Above: middle and right panels show the initial area under the curve (IAUC) from the baseline and post-OXT DCE MRI experiments overlaid onto the corresponding anatomical T2w images from one slice shown on the left panel. Below left: Placental relative signal enhancement after Dotarem^®^ injection is significantly decreased after OXT compared to baseline. Below right: Comparison of the changes in IAUC (averaged across all placentas) from baseline after either saline or OXT revealed an approximately 45% decrease after OXT, confirming that OXT-induced uterine hypercontractility was associated with significantly reduced uteroplacental perfusion (n = 3 per group). Data were analyzed with 2-tailed Student’s *t* test and presented as mean ± SEM; **p < 0.01. **B) OXT-induced hypercontractility significantly decreases placental R2*.** Above: Illustrative FSE images from an OXT-treated dam showing placental regions of interest (ROI). Below: Placental R2* was measured in 29 and 38 placentae from saline (n = 3) or OXT-treated (n = 6) dams, respectively. OXT treatment decreased placenta R2* by approximately 40% suggesting either severe uteroplacental perfusion, or profound placental squeeze draining all blood, including deoxygenated blood, away from the placenta. Data were analyzed with Mann-Whitney test and presented as mean ± SEM; ****p < 0.0001.

### OXT-induced uterine hypercontractility is associated with oxidative stress in the fetal brain

Acute uteroplacental hypoperfusion could lead to fetal hypoxemia and affect the fetal brain. Consequently, we determined the impact of OXT-induced transient placental ischemia-hypoxemia on the developing brain by assaying for lactate and oxidative stress markers (**Figure 2A-D**). Fetal brain lactate was significantly higher 4 h after OXT *vs*. saline (Figure 2A), suggesting that the impaired uteroplacental perfusion after OXT was accompanied by fetal hypoxemia. Because placental ischemia-hypoxemia induced by OXT is transient, we speculated that recovery from such an episode will be accompanied by oxidative stress. Therefore, we assayed the fetal cortex for biomarkers of oxidative stress 4 h after OXT and observed a significant increase in 4-hydroxynonenal (a lipid peroxidation product; Figure 2B) and protein carbonyl (a protein oxidation product; Figure 2C) content in pups exposed to OXT-induced hypercontractility. Concomitantly, we also detected a significant increase in the oxidized glutathione ratio in the fetal cortex after OXT suggesting a decrease in antioxidant defense (Figure 2D). These findings confirmed our prediction that acute placental ischemia from OXT-induced aberrant uterine contractility will cause oxidative stress in the fetal brain.

**Figure 2.**
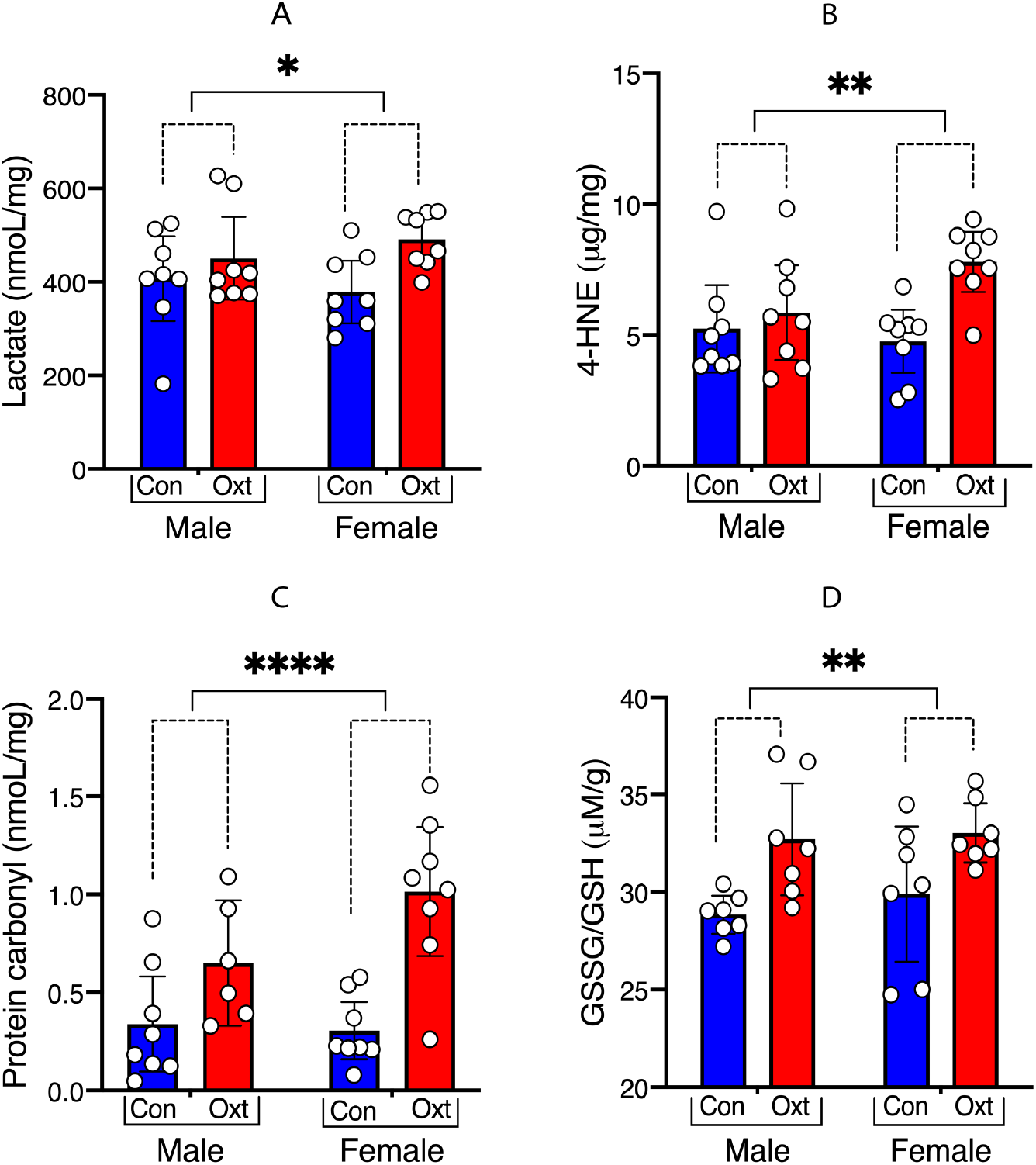
**Transient OXT-induced uterine hypercontractility causes oxidative stress in the fetal brain.** *In utero* exposure to OXT-induced uterine hypercontractility increased the concentration of lactate (Figure 2A), 4-hydroxynonenal (Figure 2B), protein carbonyl (Figure 2C), and oxidized glutathione (Figure 2D) in the fetal brain strongly suggesting the presence of oxidative stress. Data were analyzed with 2-way ANOVA and presented as mean ± SEM; *p < 0.05, **p < 0.01, ****p < 0.0001.

### Effect of OXT-induced uterine hypercontractility on the fetal cortical transcriptome

Because acute hypoxia and oxidative stress could induce transcriptomic changes in the developing brain, we assessed for this possibility by screening the fetal cortex 24 h after OXT-induced uterine hypercontractility with unbiased RNA-seq (**Figure 3A-D**). Overall, 936 of the 12660 transcripts assessed were differentially expressed (unadjusted p < 0.05) after OXT-induced hypercontractility (heatmap in Supplemental Figure 1). Notably, of the top 50 differentially expressed genes, 9 of them were mitochondrially encoded *(mt-nd2, mt-nd1, mt-nd3, mt-nd4, mt-nd4l, mt-nd5, mt-nd6, mt-cyb, mt-atp6)*. With a false discovery rate (FDR)-adjusted p ≤ 0.05, only *mt-nd2* (mitochondrially encoded NADH dehydrogenase 2 in Complex I) was significantly differentially expressed among the 936 transcripts. GO analyses revealed activation of multiple biological signaling pathways and molecular transporter activity, along with downregulation of DNA repair and nucleic acid binding activity (**Figure 3A and 3B**). KEGG pathway analyses (**Figure 3C and 3D**) revealed activation of calcium signaling mechanisms and downregulation of pathways including the Fanconi anemia pathway, important for DNA repair and known to be downregulated after hypoxic stress.^18,19^ Considering the increase in oxidative stress biomarkers, the increased expression of *mt-nd2* (essential for mitochondrial production of reactive oxygen species), and activation of calcium signaling mechanisms and the downregulation of DNA repair pathways typically observed after hypoxic injury, we concluded that one of the major effects of OXT-induced hypercontractility on the fetal brain is severe hypoxic stress. Of note, there were no significant differential expression of genes in the OXT-OXTR signaling pathway (Supplemental Figure 2).

**Figure 3.**
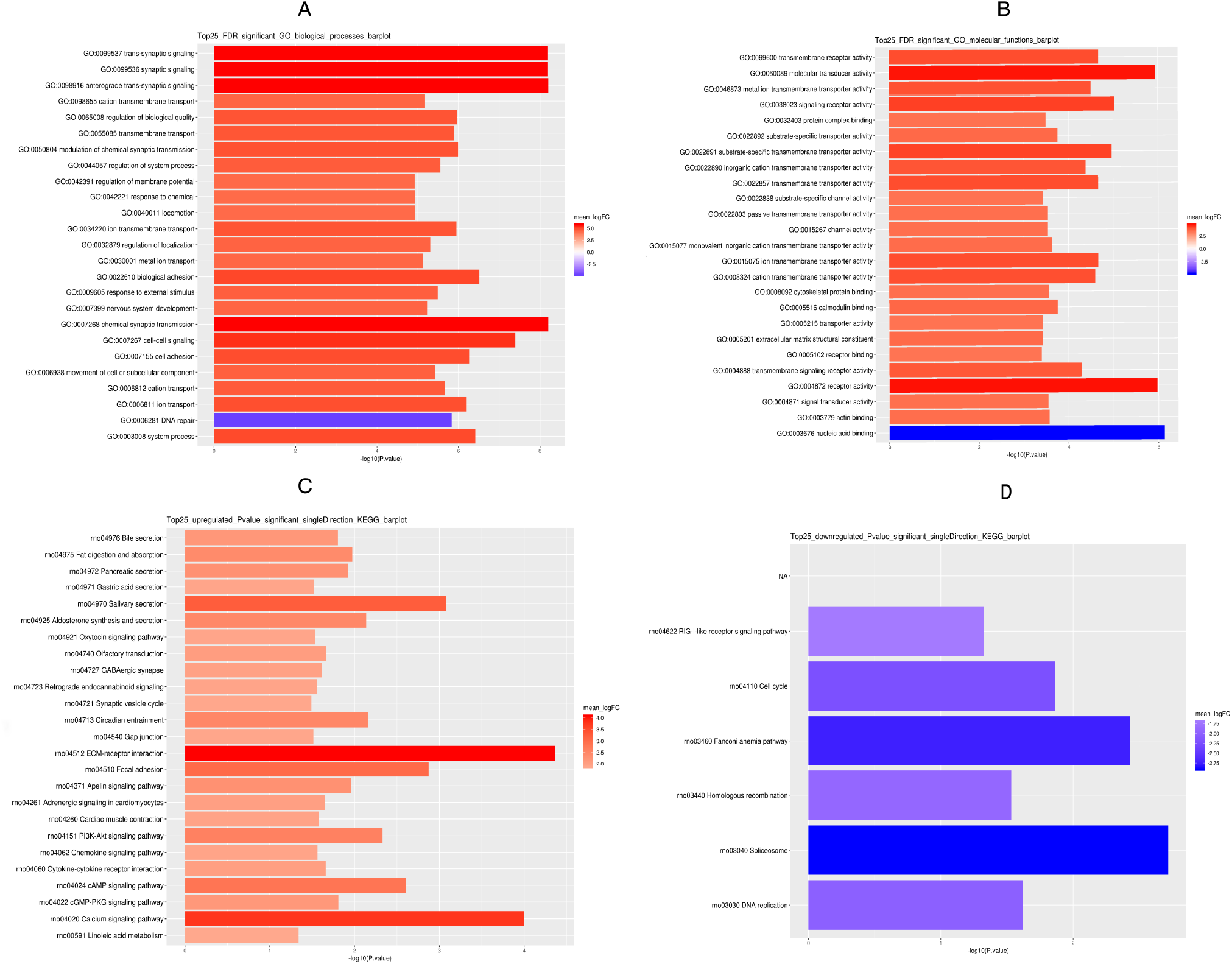
**Fetal cortical transcriptomic changes induced by OXT-induced uterine hypercontractility.** Top 25 significantly up and downregulated genes for gene ontology (GO) biological processes and molecular functions after OXT-induced hypercontractility are shown in Figures 3A and 3B, respectively. Figure 3C shows significantly upregulated genes and Figure 3D shows significantly downregulated genes with KEGG (Kyoto Encyclopedia of Genes and Genomes) analysis (n = 5 pups per group with each pup from a unique dam).

### OXT-induced uterine hypercontractility is associated with social behavioral abnormalities in the adolescent male offspring

Based on the effects of OXT-induced aberrant uterine contractility on the fetal brain, we predicted long-term functional consequences for the offspring. Therefore, in a separate set of experiments, we evaluated OXT- and saline (CON)-exposed offspring on several behavioral tests designed to assess locomotor activity, anxiety-related measures, social memory, and associative learning and memory between P28-45 (**Figure 4A-D**). There were no significant treatment or sex-related differences in general locomotor activity (1-h open-field test; **Figure 4A**) and anxiety-related measures (elevated plus maze; **Figure 4B**). However, social investigation time of a novel conspecific was significantly decreased in male, but not female, OXT-exposed offspring relative to male CON-exposed offspring (**Figure 4C**). In the observational fear learning (OFL) test (**Figure 4D**), OXT-exposed offspring exhibited significantly increased freezing levels during the last 2 min of the training period despite comparable levels during baseline and the first two minutes of training, but no sex effects were observed. However, no differences between groups were observed for the OFL contextual fear test conducted 24 h later. The differences in freezing levels between the groups during training were not likely due to differences in ambulatory activity since they performed similarly on this variable during a 1-h open-field test. In a separate set of experiments, we assessed if these behavioral changes were accompanied by differential expression of *nrxn1*, known to be associated with social behavior, in the amygdala, somatosensory cortex, and anterior cingulate cortex of P28 offspring. We detected a significant decrease in the expression of *nrxn1* in the amygdala of both male and female OXT-exposed P28 offspring, while these changes were not observed in the anterior cingulate and the somatosensory cortex (Supplemental Figure 3). Collectively, these results suggested that the main neurobehavioral effects of exposure to OXT-induced hypercontractility *in utero* are a male-specific impairment of social investigation accompanied by brain-region specific changes in the expression of genes related to social and emotional behaviors.

**Figure 4.**
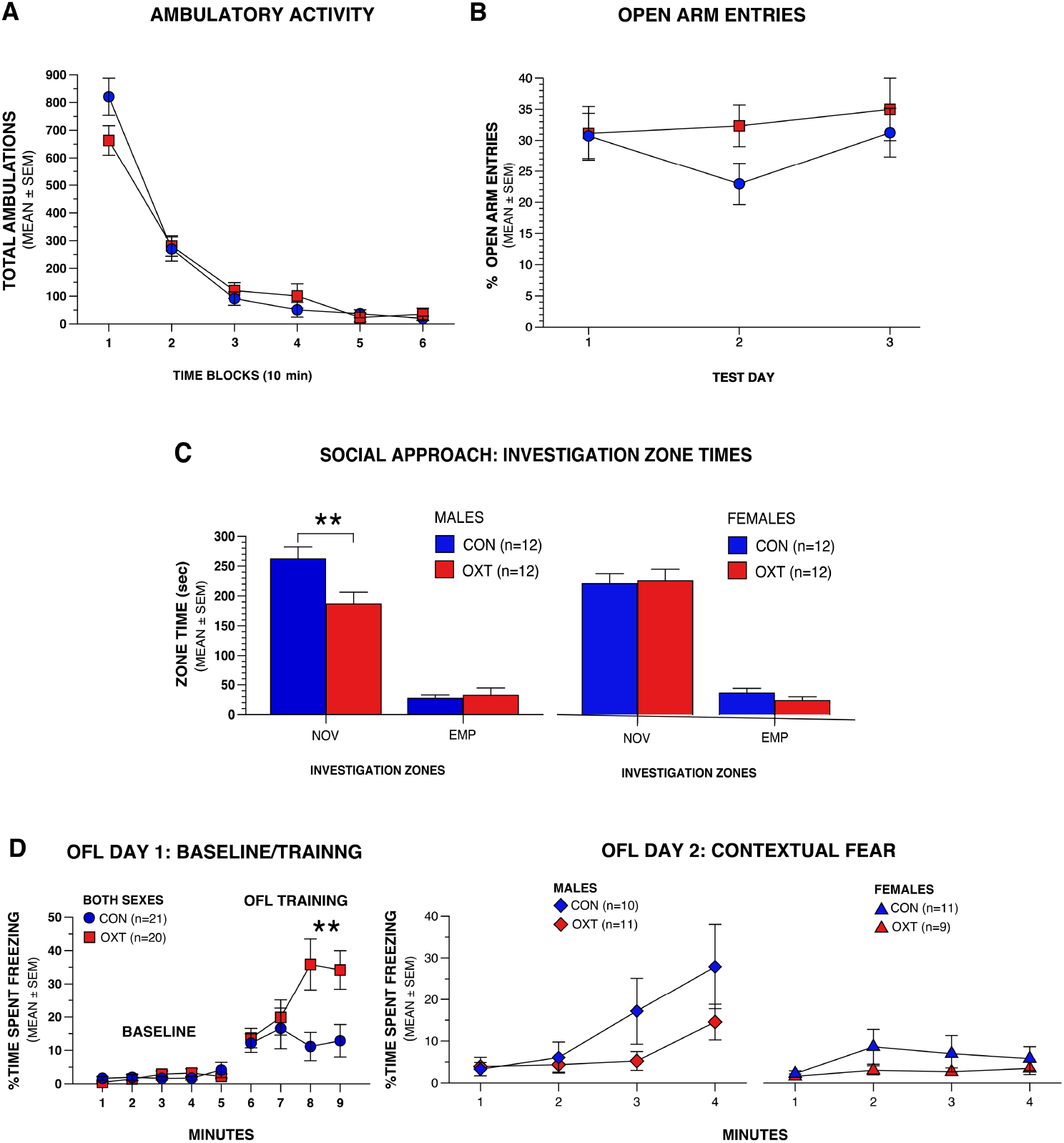
**OXT-induced uterine hypercontractility is associated with male-specific decrease in social investigation and an increase in empathy-like fear.** There were no differences in ambulatory activity in the open-field test (Figure 4A) and the percentage of open arm entries in the elevated plus maze (Figure 4B) between the CON and OXT groups. However, male OXT-exposed offspring spent significantly less time investigating a novel rat compared to CON male offspring (**p=0.007; treatment effect, *p=0.016; treatment x sex interaction, *p=0.038) (Figure 4c). No differences were observed between CON and OXT female offspring. Furthermore, OXT offspring showed significantly increased freezing levels in the OFL test during the last 2 min of training (min 8 & 9, p=0.009 for each, significant after Bonferroni correction, p<0.0125), suggesting an enhanced empathy-like response (treatment effect, *p=0.04; treatment x min interaction, **p=0.003) (Figure 4D). However, there were no differences in the contextual fear test 24 h later (n=12 male and female offspring from 9 independent litters per treatment condition).

### OXT-induced uterine hypercontractility causes persistent sex-specific increase in proteins from the mitochondrial electron transport chain

Because of sex-dependent changes in behavior, we investigated gene expression changes related to oxidative stress 1 h after OXT-induced uterine hypercontractility in both male and female offspring. Specifically, we performed planned comparisons of treatment-related differences in the expression of select target genes related to ETC/oxidative stress, antioxidant defense, and hypoxia/apoptosis pathways using prevalidated Taqman^®^-specific probes. We attempted to perform this in the anterior cingulate cortex and amygdala, but quick liquefaction of the E21 pup brain prevented us from accurate dissection of these brain regions. Therefore, the entire fetal cortex was assessed. Multiple genes were differentially expressed in a sex-dependent manner; overall, there was an increased expression of genes related to ETC/oxidative stress *(mt-nd4, mt-nd5, mt-atp8)*, anti-oxidant defense *(sod2, cygb)*, and hypoxia/apoptosis *(casp8, hif1a)* pathways in treated males vs. females (**Figure 5A**). *Oxtr* expression was not significantly different with OXT nor was there a sex x OXT interaction at this timepoint (Supplemental Figure 4). We then selectively quantified changes in proteins emblematic of these pathways with immunoblots. HIF-1a1pha and cleaved caspase-3 (CC3) were assessed in nuclear and whole cortical lysates, respectively, at 1 h and 18 h. There were no treatment or sex-related differences in either HIF-1a1pha or CC3 (Supplemental Figure 5), suggesting that the related gene expression changes were probably transient. Because of the predominance of mitochondrially coded genes among genes that were differentially expressed at 24 h after OXT-induced hypercontractility, we focused on OXPHOS (oxidative phosphorylation) proteins in the anterior cingulate cortex, considering the importance of this brain region for social behavior and cognition.^20–25^ OXPHOS proteins were quantified in the mitochondrial fraction isolated from the anterior cingulate cortex of male and female P28 offspring. We observed that ETC complex I, III, and IV proteins were selectively higher in the anterior cingulate cortex of P28 male OXT-exposed offspring (**Figure 5B**), suggesting that transient but severe hypoxic events *in utero* can cause persistent mitochondrial changes.

**Figure 5.**
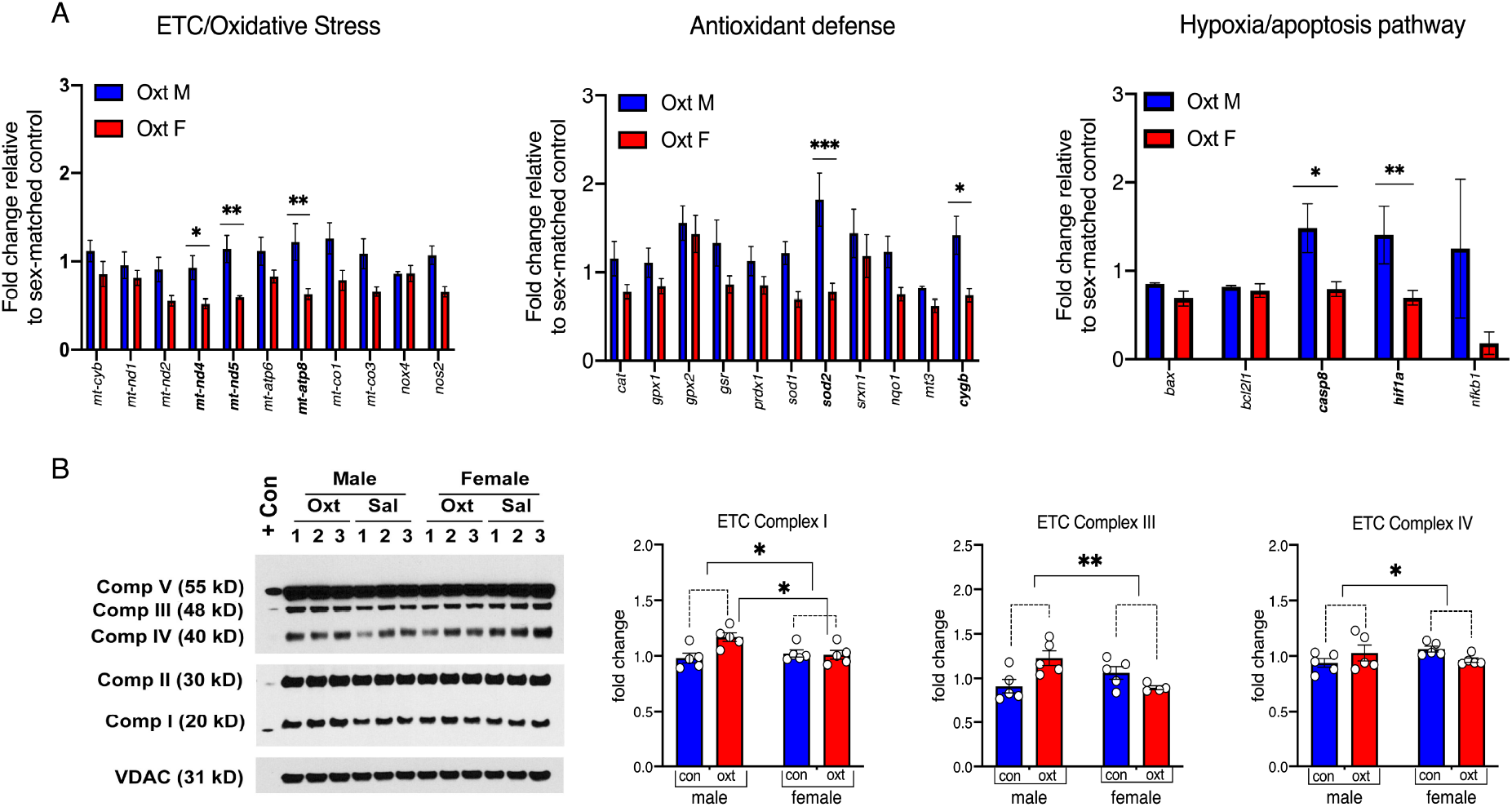
**A) OXT-induced hypercontractility causes sex-dependent gene expression changes in the E21 fetal cerebral cortex.** Multiple genes related to ETC/oxidative stress *(mt-nd4, mt-nd5, mt-atp8*), antioxidant defense *(sod2, cygb)*, and hypoxia/apoptosis *(casp8, hif1a)* pathways were differentially expressed in a sex-dependent manner 1 h after OXT-induced hypercontractility. Overall, there was an increased expression of genes in treated males vs. females. *p<0.05, **p<0.01, and ***p<0.001. **B) OXT-induced hypercontractility causes enduring upregulation of OXPHOS proteins in the anterior cingulate cortex of male P28 offspring.** Left: Representative immunoblots of anterior cingulate cortex homogenates from P28 offspring showing increased expression of proteins related to the mitochondrial electron transport chain (OXPHOS) in males. Scatter plots show a significant treatment x sex interaction for complex I (*p=0.02; [F (1, 16) = 6.5]), complex III (**p=0.004; [F (1, 15) =12.0]), and complex IV (*p=0.04 for treatment x sex interaction, [F (1, 16) = 4.6]). Normalization was done with mitochondrial VDAC1 protein. Data were analyzed with 2-way ANOVA and presented as mean ± SEM; *p < 0.05, **p < 0.01, ***p < 0.001.

### OXT-exposed male offspring demonstrate enduring impairment in cortical functional connectivity

Complex behavioral changes cannot be explained by differences in gene and protein expression alone. Therefore, we examined the effects of OXT exposure on systems-level brain organization using functional connectivity optical intrinsic signal imaging (fcOISI)^26^ in male offspring exposed to either OXT or saline. Candidate regions of interest (ROIs) were placed in cortical areas corresponding to cingulate, motor, somatosensory, retrosplenial and visual networks (**Figure 6A**). Saline-treated rats exhibited robust patterns of functional connectivity that feature prominently in healthy rats. These characteristics included strong positive (reds) ipsilateral correlation adjacent to each ROI, as well as mirrored homotopic functional connectivity contralateral to each ROI (**Figure 6B**). Additionally, strong anticorrelations (blues) were present between opposed functional networks, e.g. amongst sensorimotor and retrosplenial cortices. These features were altered in the OXT-treated group. For example, correlation difference maps reveal significantly reduced homotopic functional connectivity strength in cingulate cortex (**Figure 6C**), along with large changes in ipsilateral connectivity within cingulate, visual and retrosplenial regions (Reds, **Figure 6C**). The magnitude of anticorrelations between sensorimotor and visual and parietal regions, as well as between retrosplenial and lateral sensory regions was also attenuated (Blues, **Figure 6C**). One of the most salient differences between saline and OXT-related rats was the spatial extent of positive correlations exhibited by a given ROI. To quantify this metric, we constructed maps of global node degree, calculated as the number of cortical functional connections exhibited by a given pixel having a correlation strength higher than z(r)>0.3 (**Figure 6D**). Saline-treated animals exhibited high connection number (yellows, whites) in retrosplenial, visual and motor regions. OXT-treated rats exhibited more global reductions in node degree, as indicated by the darker shades of red and black over large portions of the cerebral cortex. Significant regional differences in this measure were observed in portions of motor, somatosensory, parietal, cingulate, and visual cortices (**Figure 6D**). Taken together, exposure to OXT-induced uterine hypercontractility *in utero* affects functional brain organization within and across functionally distinct brain systems.

**Figure 6.**
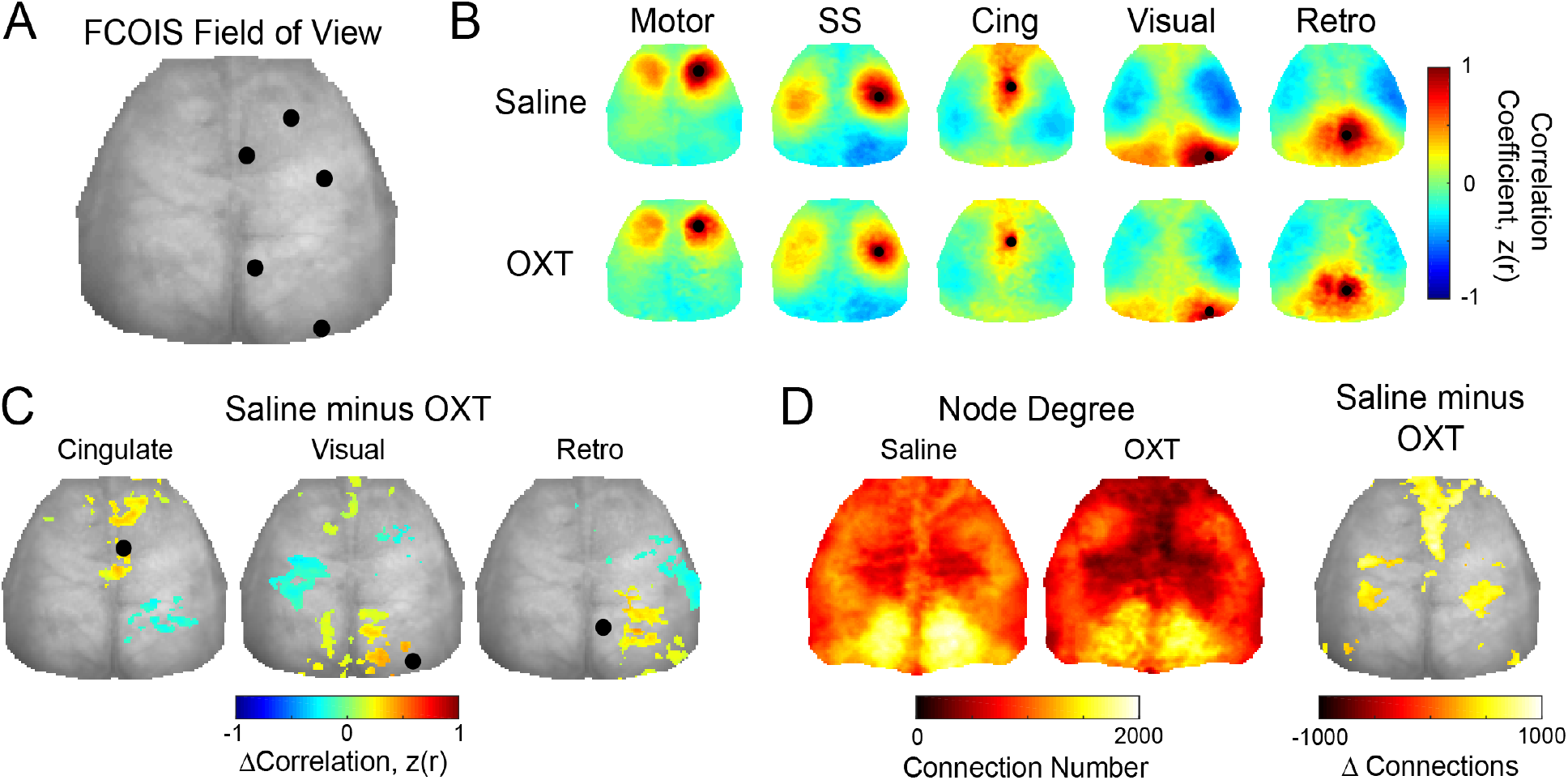
Functional connectivity with optical intrinsic signaling. ***In utero* OXT exposure affects functional brain organization in juvenile male offspring.** Functional connectivity optical intrinsic signal imaging was performed in P23 male offspring exposed to either OXT (n=5) or saline (n=7) *in utero*. **A) FCOIS field of view.** Seeds for functional connectivity analysis (black dots on representative white light image of the rat skull) were placed in cortical areas corresponding to cingulate, motor, somatosensory, retrosplenial and visual networks. **B) Top Row:** Saline-treated rats exhibited patterns of functional connectivity with strong positive (reds) ipsilateral correlation adjacent to and contralateral from each seed and strong anticorrelations (blues) between opposed functional networks (e.g. between sensorimotor and retrosplenial cortices). Qualitatively, these general features of healthy patterns of functional connectivity were reduced in OXT-treated rats (Bottom Row). **C) Functional connectivity difference maps in select networks.** Significant reduction in ipsilateral connectivity was observed in cingulate, visual, and retrosplenial regions. Anticorrelated activity (blues) was also significantly reduced between somatomotor areas and visual regions, along with lateral sensory and retrosplenial regions. **D)** Maps of node degree for each group were calculated by thresholding each pixel’s functional connectivity map at z(r)>0.3 and counting all pixels above this threshold. Saline-treated animals exhibited high connection number (yellows, whites) in retrosplenial, visual and motor regions. OXT-treated rats exhibited more global reductions in node degree. Significant regional differences in node degree were observed in portions of motor, somatosensory, parietal, cingulate, and visual cortices (difference map). Significance determined by 2-tailed Students t-test assuming unequal group variance at each pixel and corrected for multiple comparisons using FDR correction on a cluster-wise basis.

## Discussion

In this translational study with term pregnant rats, we show that OXT-induced aberrant uterine contractility was associated with severe but transient impairment of uteroplacental perfusion with significant functional consequences for the offspring. Specifically, we observed significant oxidative stress in the fetal brain, with an increase in both biomarkers and genes mediating oxidative stress. These changes were accompanied by social behavioral abnormalities especially in males, significant reductions in functional connectivity between different brain regions, and an enduring upregulation of mitochondrial ETC complexes in the anterior cingulate cortex. Collectively, our findings suggest that even transient but severe impairment of placental perfusion can have lasting detrimental consequences for the offspring.

Our studies appear to be the first to comprehensively examine the impact of transient but severe intrapartum hypoxemia resulting from aberrant uterine contractility on the fetal brain. Our model is more pragmatic in that it avoids complex vascular surgery to reduce uteroplacental perfusion, but more importantly, allows for quick reversibility of the insult to better reflect complications during human labor.^13,14,27–29^ Our findings challenge the prevailing assumption that neonatal metabolic status is a good surrogate for the wellbeing of the fetal brain and suggests that even a short duration hypoxic insult could be damaging to the fetal brain. Furthermore, our findings offer mechanistic evidence linking intrapartum hypoxia, oxidative stress, and persistent abnormalities in functional connectivity as the possible drivers of neurodevelopmental disorders observed after acute obstetric complications.^1–5,30–34^

Disruption to the OXT-OXTR system could possibly play a role in the observed biochemical and behavioral abnormalities. We minimized this possibility by choosing a dose of OXT that was not associated with significant placental transfer in our model.^35^ Though we cannot definitively rule out activation of transient OXTR signaling at the time of OXT injection, multiple lines of evidence from our studies suggested minimal involvement of the OXT-OXTR system (RNA-seq data, Taqman qPCR) at a later time point (Supplemental Figure 2). This may appear be at odds with the study by Kenkel et al. showing acute effects of OXT on the fetus,^36^ but we posit a few possibilities why our outcomes were different: (i) we administered OXT intravenously causing a profound and immediate tetanic uterine contraction, compared to an intraperitoneal injection in the Kenkel study, and (ii) the associated drastic reduction in uteroplacental perfusion most likely decreased the delivery of OXT-containing maternal blood to the placenta and to the fetus. Our study is perhaps more comparable to the study by Boksa et al. demonstrating that acute administration of OXT before birth worsened anoxic brain injury in the neonatal rat brain.^37^ With this background, we believe that the observed effects in the developing brain are more likely due to indirect effects from the tetanic uterine contraction rather than the direct effects of OXT.

Observation of severe reduction in uteroplacental perfusion after OXT-induced uterine contractility motivated us to explore hypoxic stress in the fetal brain as a possibility. This line of investigation was reinforced by evidence for increased fetal brain lactate and biomarkers of oxidative stress after OXT. Though examination of the global fetal cortical transcriptomic changes 24 h after OXT did not reveal involvement of specific hypoxia-associated pathways, there was a preponderant differential expression of genes from the mitochondrial electron transport chain. Taken together with the central role of brain mitochondria in responding to hypoxic and oxidative stress,^38–41^ and the brain-region specific persistent alteration of mitochondrial ETC proteins in juvenile offspring, we were convinced that one of the major effects of OXT-induced uteroplacental ischemia was oxidative stress with possible mitochondrial dysfunction.

A major finding of our behavioral studies was an impairment of social investigation in male offspring exposed to OXT-induced hypercontractility *in utero*. We pursued this with evaluation of the expression of *nrxn1*, previously associated with social behavior, in brain regions known to be integral to the neurocircuitry of social interaction (amygdala, anterior cingulate cortex, somatosensory cortex).^42–49^ Expression of *nrxn1* was differentially regulated in the brain regions of OXT-exposed offspring regardless of the sex; *nrxn1* was significantly downregulated in the amygdala, but unchanged in the anterior cingulate and the somatosensory cortices of OXT vs. control offspring. Because these results do not explain male-specific impairment of social behavior, we speculate that these changes could be due to sex differences in the response to oxidative stress.^38,50–54^ Supporting this viewpoint is the significant sex difference in the expression of genes mediating oxidative stress 1 h after OXT, and the presence of persistently altered mitochondrial ETC proteins in the anterior cingulate cortex of only P28 male offspring.

Mitochondrial dysfunction has also been recently demonstrated to lead to significant reductions in long range axonal connectivity.^55^ Because structural connectivity is a required substrate for functional connectivity, OXT-induced changes in factors affecting anatomical connections between neurons could affect functional brain network organization (i.e., segregation and integration). Our general finding of impaired cortical functional connectivity in male offspring treated with OXT is broadly supportive of this notion. More specifically, OXT-treated rats exhibited significant reductions in the magnitude of anticorrelations between functionally distinct networks positioned across the anterior-posterior axis. In humans, observations of anticorrelated spontaneous brain activity resulted in some of the first accounts of intrinsic brain organization.^56^ Dynamic patterns of anticorrelated activity segregate distinct processes across networks during both task and resting state paradigms, ^56,57^ with the most common example in humans occurring between the dorsal attention network and default mode network^58^. Altered synchrony across large cortical distances, as observed by reduced anticorrelated activity, could prevent inter-network communication necessary for higher-order integration. Another marked distinction between saline- and OXT-treated rats was significantly altered ipsilateral node degree in portions of motor, somatosensory, parietal, cingulate, and visual cortices. These reductions could be indicative of network-level inefficiencies important for segregation. For example, in mice we have recently shown^59^ that improved tactile proprioception after stroke was associated with increased node degree in motor, somatosensory, and parietal cortices – all regions relevant for processing proprioception and touch.^60^ In the present study, reduced within-network connectivity (i.e., node degree) combined with poorer integration across networks (i.e. reduced anticorrelations) could impair multi-domain behavioral output, as demonstrated by poorer social and emotional performance of the OXT-treated group.

Our findings have to be interpreted in the context of a few limitations. First, it is not possible to extrapolate the duration of placental ischemia-hypoxia in rats to humans. Furthermore, human babies are likely to be delivered if hypoxia persists long enough to cause concerning fetal heart rate changes. Nevertheless, the developing human brain is generally more sensitive to hypoxia than the rodent brain and is more likely to be susceptible to mild hypoxic injury.^61^ Second, we did not treat placental ischemia-hypoxia with oxygen administration in the dam, a practice that is common during human labor. Therefore, there is a possibility that oxidative stress could be worsened in this setting. Third, we focused only on 3 brain regions based on behavioral changes, but it is likely that we may have missed important changes elsewhere. Similarly, a brain-region specific analysis of oxidative stress immediately after OXT was not possible because easy liquefaction of the fetal brain prevented an accurate dissection of brain regions. Despite these limitations, our findings provide detailed insight into the possible mechanisms by which transient placental ischemia-hypoxia could induce persistent dysfunction in the developing brain.

In conclusion, transient intrauterine ischemia-hypoxemia was associated with male-specific differences in social behavior, differential expression of selective oxidative stress genes in the developing cortex, and enduring upregulation of mitochondrial ETC proteins in the anterior cingulate cortex of male offspring. Collectively, our findings support the idea that severe but reversible hypoxic stress could induce permanent dysfunction in the developing brain and raises the possibility of persistent mitochondrial dysfunction as the link between transient intrauterine asphyxia and neurodevelopmental disorders.

## Methods

### 1. Animals and Drugs

Timed pregnant Sprague Dawley (SD) rats (Charles River Laboratories, Wilmington, MA) were used for all experiments. Oxytocin (OXT; Selleck Chemicals, Houston, TX) was prepared as a 1mg/mL solution in sterile normal saline and diluted as necessary. We performed a pilot study with 3 different doses of OXT (10 mcg/kg, 100 mcg/kg, and 1 mg/kg) to determine the duration of externally visible uterine contraction and confirmed that 100 mcg/kg caused a predictable 10-15 min tetanic uterine contraction. Uterine contraction induced by 10 mcg/kg, however, was variable in length, while that induced by 1 mg/kg was so severe that it occasionally resulted in stillborn pups. Furthermore, 100 mcg/kg OXT administered to the mother was not accompanied by an increase in fetal brain OXT in our previous study,^35^ suggesting a lack of placental transfer at this dose. During the course of these experiments, we also developed a rapid method to identify the sex of the pups using visual inspection of the anogenital distance at E21. Another experimenter cross-checked and verified the sex by performing a minilaparotomy to identify the presence (male) or absence (female) of testes and seminal vesicles. The total number of animals used for specific experiments is detailed in Supplemental Table 1. Technical details are outlined in Supplemental Tables 2 and 3. Litter data and weight gain trajectories of the offspring are presented in Supplemental Figure 6.

### 2. Placental imaging

For the dynamic contrast enhanced (DCE) MRI scans, SD dams at E21 (n=3) were anesthetized with isoflurane in 100% oxygen and imaged on an Agilent (Santa Clara, CA) 4.7T DirectDrive MRI system using a 7.2 cm inner diameter quadrature RF coil. Anatomic scans were initially acquired across the entire abdomen using a respiratory-gated T2-weighted 2D multi-slice fast spin echo sequence (FSEMS; 0.5 x 1 x 1 mm^3^ resolution, zero-padded to 0.5 x 0.5 x 1 mm^3^, TR/TE = 2000/48 ms, echo train length = 4, 4 averages, respiratory gated, imaging time approximately 6 min depending upon respiration rate). Placental blood flow was measured by DCE MRI with a tail vein injection of Dotarem^®^ (Guerbet, Princeton, NJ, 0.2 mmol/kg). The rats were then injected with 100 μg/kg of OXT through a 22G tail vein catheter to induce tetanic uterine contraction. Following the cessation of gross abdominal motion, typically around 30 min as assessed by non-gated low-resolution scouts, the anatomic scan was re-acquired to account for placental displacement during the contraction period and the DCE measurement was repeated with a repeat dose of the same contrast agent. Placental perfusion was quantified as the initial area under the curve (IAUC) of the DCE time-intensity for each placental voxel using the DCE@urLAB package.^62^ Each rat placenta was manually segmented from pre- and post-OXT anatomical images using customized MATLAB routines and interpolated onto the IAUC maps. As the DCE protocol involved two injections of contrast agent separated by approximately 50 min, the time under anesthesia and the presence of residual contrast agent from the first injection could potentially distort the post-OXT DCE results. To examine the size of these effects, we repeated the DCE protocol where we injected the same volume of saline instead of OXT (n=3) followed by comparison of the % change in IAUC between the saline and OXT groups. Placental R2* mapping was performed using a 3D multi-echo gradient echo sequence (0.75 mm isotropic resolution, TR/TE/ΔTE = 40/1.7/4 ms, 6 echoes, flip angle = 15 degrees, 4 min 16 sec) to measure changes in deoxyhemoglobin concentration after 100 μg/kg of OXT (n=6) or an equivalent volume of saline (n=3) in a separate set of experiments. Rat placentas were manually segmented from pre- and post-OXT images using customized MATLAB routines, and optimal estimates of R2* were determined for each placenta using software implementing Bayesian probability theory. For the saline-injected rats, a 30-min delay was observed prior to acquiring post-injection scans.

### 3. Lactate assay

To determine if OXT-induced uterine contractions induce hypoxemia in the fetal brain, we assayed for fetal brain lactate 4 h after 100 μg/kg i.v OXT or saline in timed pregnant E21 Sprague Dawley (SD) dams (n = 8 per group). Briefly, whole male and female fetal cerebral cortices were homogenized on ice and deproteinized with 10 kDa molecular weight cut-off spin filter to remove lactate dehydrogenase. Protein concentrations were determined using BCA Protein Assay Kit (ThermoFisher Scientific), followed by lactate assay (Lactate Colorimetric Assay Kit II; Sigma Aldrich) according to manufacturer’s instructions. Results calculated from the standard curve were expressed as nmoL/mg brain protein.

### 4. Oxidative stress assays

To determine if OXT causes oxidative stress in the fetal brain, we assayed male and female fetal brains for 4-hydroxynonenal (4-HNE, a byproduct of lipid peroxidation), protein carbonyl, and the antioxidant GSSG/GSH from the same E21 dams used for the lactate assay. Briefly, whole fetal cerebral cortices from male and female pups were homogenized in PBS buffer on ice. Protein concentrations were determined using BCA Protein Assay Kit (Thermo Scientific), followed by 4-HNE assay (OxiSelect™ HNE Adduct Competitive ELISA Kit), protein carbonyl (OxiSelect™ Protein Carbonyl ELISA kit), and total glutathione assay (OxiSelect™ GSSG/GSH Assay Kit) according to manufacturer’s instructions. All assays were supplied by Cell Biolabs, Inc, San Diego, CA. Results calculated from the standard curve were expressed as mcg/mg of brain protein.

### 5. RNA-seq and analysis

E21 timed pregnant SD rats were treated with either 100 μg/kg OXT or saline through a 22G tail vein catheter (n = 5 each). 24 h later, fetal cerebral cortices were harvested under isoflurane anesthesia. Total RNA was extracted the right cerebral cortex using RNAeasy kit (Qiagen). Only RNA with RIN > 9.5 were used for RNA-seq. Samples were prepared according to library kit manufacturer’s protocol, indexed, pooled, and sequenced on an Illumina HiSeq. Basecalls and demultiplexing were performed with Illumina’s bcl2fastq software and a custom python demultiplexing program with a maximum of one mismatch in the indexing read. RNA-seq reads were then aligned to the Rattus norvegicus Ensembl Rnor_5.0 top-level assembly with STAR version 2.0.4b. Gene counts were derived from the number of uniquely aligned unambiguous reads by Subread: featureCount version 1.4.5. Isoform expression of known Ensembl transcripts were estimated with Sailfish version 0.6.13. Sequencing performance was assessed for the total number of aligned reads, total number of uniquely aligned reads, and features detected. The ribosomal fraction, known junction saturation, and read distribution over known gene models were quantified with RSeQC version 2.3. All gene counts were then imported into the R/Bioconductor package EdgeR and TMM normalization size factors were calculated to adjust for samples for differences in library size. Ribosomal genes and genes not expressed in at least 4 samples greater than one count-per-million were excluded from further analysis. The TMM size factors and the matrix of counts were then imported into the R/Bioconductor package Limma. Unknown latent effects were then estimated with the R/Bioconductor package SVA. Weighted likelihoods based on the observed mean-variance relationship of every gene and sample were then calculated for all samples with the voomWithQualityWeights. The performance of all genes was assessed with plots of the residual standard deviation of every gene to their average log-count with a robustly fitted trend line of the residuals. Differential expression analysis was then performed to analyze for differences between conditions and the results were filtered for only those genes with Benjamini-Hochberg false-discovery rate (FDR) adjusted p-values ≤ 0.05. For each contrast extracted with Limma, global perturbations in known KEGG pathways was detected using the R/Bioconductor package GAGE9 to test for changes in expression of the reported log 2-fold-changes reported by Limma in each term versus the background log 2 fold-changes of all genes found outside the respective term. The R/Bioconductor package Pathview10 was used to display heatmaps or annotated KEGG graphs across groups of samples for each KEGG pathway with a p-value less than or equal to 0.05. RNA-seq data have been deposited in NCBI’s Gene Expression Omnibus and are accessible through GEO Series accession number GSE146155 (https://www.ncbi.nlm.nih.gov/geo/query/acc.cgi?acc=GSE146155).

### 6. Behavioral experiments

Male and female offspring of E21 SD dams (n = 9 each) treated with OXT (100 μg/kg) or saline (OXY or CON, respectively) were subjected to behavioral testing. There were no differences in litter size, sex ratio, or survival of the offspring from both treatment groups. Pups were weaned on P21 and subsequently evaluated on a battery of behavioral tests from postnatal day (P) 26-45 in the following order: 1) 1 h open field; 2) social approach; 3) elevated plus maze; and 4) observational fear learning. The OXT and CON rats were tested in two cohorts (n = 6 and 3 dams each, respectively) for which identical behavioral protocols were used. A total of 12 male and female offspring from the OXT and CON group were tested. Details of all behavioral experimental procedures are described previously, ^63^ and included in the Supplemental File.

### 7. Taqman qPCR

Male and female fetal brains exposed to either 100 μg/kg OXT or saline (n= 8 per group) *in utero* were harvested and immediately stored at −80°C. RNA was extracted using RNeasy plus mini kit (Qiagen), converted to cDNA using SuperScript^®^ IV VILO™ master mix kit (Invitrogen), and 25 ng of the template cDNA was then combined with a ready-to-use TaqMan Fast Advanced qPCR Master Mix (Thermo Fisher Scientific, Waltham, MA) for experiments in a pathway-specific custom TaqMan array (Custom TaqMan^®^ Array Fast Plate 96, Life Technologies, Carlsbad, CA). Expression levels of 28 genes relevant to three specific pathways mitochondrial electron transport chain (ETC) complexes *(mt-cyb, mt-nd1, mt-nd2, mt-nd4, mt-nd5, mt-atp6, mt-atp8, mt-co1, mt-co3, nox4, nos2)*, oxidative stress and anti-oxidant defense *(cat, gpx1, gpx2, gsr, prdx1, sod1, sod2, srxn1, nqo1, mt3, cygb)*, and hypoxia, stress and toxicity *(bax, bcl2l1, casp8, hif1a, nfkb1)* were assayed in triplicate along with four endogenous housekeeping control genes *(18S rRNA, gapdh, pgk1*, and *actb)*. To better understand the behavioral differences in P28 offspring, we examined the expression of *nrxn1* in the amygdala, anterior cingulate cortex, and the somatosensory cortex of male and female P28 offspring in a separate set of experiments. Thermal cycling was performed in 7500 Fast Real-Time PCR System (Applied Biosystems^®^, Foster City, CA) and the threshold cycle (Ct) values for all genes were calculated using proprietary software. We tested the experimental stability of all 4 endogenous reference genes using the geNorm algorithm and determined *pgk1* to be the most stable reference gene for oxidative stress studies, and both *pgk1* and *actb* for the learning-associated studies. Relative mRNA expression, normalized either to *pgk1* or a geometric mean of *pgk1* and *actb*, was calculated using the 2^-ΔΔCT^ method with sex-matched control samples as reference.

### 8. Western blotting

Lysates for western blot were prepared from approximately 100 mg of fetal cortex using RIPA buffer (50 mM Tris HCl pH 7.5, 150 mM NaCl, 2 mM EDTA, 1% NP40, 0.1% SDS) with protease and phosphatase inhibitor cocktail (ThermoFisher Scientific Inc). Mitochondrial and nuclear fractions from the fetal cortex were separately prepared using appropriate kits (mitochondrial isolation kit, catalog # 89801 and subcellular protein fractionation kit, catalog # 87790, ThermoFisher Scientific Inc) following manufacturer’s instructions. Approximately 30 μg of protein was subjected to gel electrophoresis and transferred to membrane using Bolt western blot reagents from ThermoFisher Scientific Inc (bolt 4-12% Bis Tris gel, catalog # NW04125; bolt sample reducing agent, catalog # B0009; bolt LDS sample buffer, catalog # B0007; iBlot2 dry blotting system). Membranes were blocked with TBST buffer (catalog # S1012, EZ Bioresearch), containing 5% milk for 1 hour at room temperature on a shaker. Following a brief wash with TBST buffer, the membranes were immunoblotted overnight at 4 °C on a shaker with antibodies against hif1a (Catalog # 14179, Cell Signaling Technology), cleaved caspase 3 (Asp-175) (Catalog # 9661, Cell Signaling Technology), OXPHOS rodent ab cocktail (ab110413) (Catalog # MS604300, Abcam Inc). Beta-actin (Catalog # MA5-11869, ThermoFisher Scientific Inc), VDAC1 (Catalog # ab 15895, Abcam Inc), and histone H3 (Catalog # 9715S, Cell Signaling Technology) were used for normalization. For OXPHOS experiments, rat heart mitochondria were used as positive control. All the primary antibodies were used at a dilution of 1:1000. HRP-conjugated secondary antibodies (anti-Rab IgG, catalog #7074 and anti-mouse IgG, catalog #7076, Cell Signaling Technology) were used at a dilution of 1:1000 for 1 hour at room temperature on a shaker. Immunoblots were incubated with Western ECL substrate (catalog # 1705060, Bio-Rad Inc) for 5 minutes at room temperature, followed by exposure to film inside a cassette in dark room, and developed using Konica Minolta Inc, Film Processor (catalog # SRX-101A). Western blot images were processed with ImageJ for densitometric quantification.

### 9. Functional connectivity analysis

Functional connectivity of the brain of OXT and saline-exposed male offspring was assessed at P23 using a reflectance-mode functional connectivity optical intrinsic signal (fcOIS) imaging system.^64^ The fcOIS system images the brain through the intact skull and records spontaneous fluctuations in oxy- and deoxyhemoglobin. Through neurovascular coupling, these fluctuations represent spatio-temporal variations in neural activity, similar to human BOLD-fMRI. Briefly, juvenile rats were anesthetized with an i.p injection of ketamine (86.9 mg/kg) and xylazine (13.4 mg/kg) as described previously.^65^ Optical intrinsic signals were acquired during sequential elimination provided by four polarized light emitting diodes (LEDs, Thorlabs, 470 nm: M470L3-C1, 590 nm: M590L3-C1, 617 nm: M617L3-C1, and a 625 nm: M625L3-C1) placed approximately 20 cm above the rat’s head. Diffuse reflected light from the rat head was detected by a cooled, frame-transfer EMCCD camera (iXon 897 Ultra, Andor Technologies). A crossed linear polarizer was placed in front of the camera lens to reject specular reflection from the LEDs. The LEDs and camera were time-synchronized and controlled via a National Instruments data Acquisition card (NI PCI 6733) and computer (Dell Workstation) using custom-written software (MATLAB, Mathworks) at a frame rate of 120 Hz (30 Hz/LED). Image processing and functional connectivity measurements (including seed-based functional connectivity, homotopic functional connectivity, and regional node degree) were performed as described previously.^59^ Group averaged correlation maps and matrices are presented in Supplemental Figure 7.

### 10. Statistics

For assay and biochemical data, outliers were detected and eliminated using ROUT (robust regression and outlier analysis) with Q set to 10%. Normality of residuals was checked with D’Agostino-Pearson omnibus test. Data with non-normal residuals were Box-Cox transformed prior to statistical testing. For experiments where sex was not included as a variable (i.e., placental imaging), data were analyzed with student’s t-test. For experiments where sex of the offspring was included as a variable, data were analyzed with a 2-way ANOVA followed by Sidak’s multiple comparisons test. RNA-seq data were analyzed with generalized linear models to test for gene/transcript level differential expression. Differentially expressed genes and transcripts were then filtered for false discovery rate (FDR) adjusted p-values ≤ 0.05. Behavioral data were analyzed with repeated-measures ANOVA models to evaluate effects of treatment, sex, and time-related variables. Following a statistically significant interaction between the main factors, simple main effects were calculated to provide clarification of statistically significant between-treatment and within-treatment differences. Where appropriate, the Huynh-Feldt adjustment was used to protect against violations of sphericity, and Bonferroni correction was applied to multiple pairwise comparisons. Prior to statistical testing of fcOIS measures, Pearson r values were transformed to Fisher z-scores followed by a two-tailed student’s t-test. Seed-based comparisons across groups were performed using a two-tailed student’s t-test assuming unequal group variance, followed by FDR correction for multiple comparisons. Maps of node degree were compared at each pixel using a two-tailed student’s t-test assuming unequal group variance and corrected for multiple comparisons using clustering statistics. Differences between groups were considered statistically significant if p < 0.05 following correction for multiple comparisons. Data are presented as mean ± SEM. Statistical analyses of biochemical data were performed with Prism 8 for macOS (version 8.2.1; GraphPad Software Inc.), behavioral data with SYSTAT12 (Chicago, IL), and fcOISI data with MATLAB.

### 11. Study approval

All experiments reported here were approved by the Institutional Animal Care and Use Committee at Washington University in St. Louis (#20170010) and comply with the ARRIVE guidelines.

## Author contributions

AP conceived the study design, planned the experiments, and wrote the manuscript with help from all co-authors. AP, TG and JJ developed the OXT dosing paradigm and conducted the molecular biology experiments. JDQ and JRG performed the placental imaging experiments, analyzed the data, and helped draft the manuscript. SBC performed the behavioral experiments, and SEM and DFW conducted the analysis and helped with interpretation and drafting of the manuscript. AB and AQB collected, analyzed, and interpreted the fcOISI data, and helped write the manuscript. NR provided the clinical context and assisted with drafting of the final manuscript.

## Acknowledgments

This work was supported by startup funds from the Department of Anesthesiology, Washington School of Medicine (AP) and NIH R01 grant NS102870 (AQB). Placental imaging studies presented in this work were carried out, in part, using the Small Animal MR Facility of the Mallinckrodt Institute of Radiology, Washington University in St. Louis. Animal behavioral studies were conducted by the Animal Behavior Core, a subcore of the Washington University Intellectual and Developmental Disabilities Research Center. Genomic analysis was conducted at the Genome Technology Access Center (GTAC) in the Department of Genetics at Washington University School of Medicine. GTAC is partially supported by NCI Cancer Center Support Grant #P30 CA91842 to the Siteman Cancer Center and by ICTS/CTSA Grant# UL1TR002345 from the National Center for Research Resources (NCRR), a component of the National Institutes of Health (NIH), and NIH Roadmap for Medical Research.

