## Supplemental File for "Transient uterine hypercontractility causes fetal cerebral oxidative stress and enduring mitochondrial and behavioral abnormalities in adolescent male rat offspring"

### 1. Supplemental Tables

**Table 1:** Details of animal use

| Experiment | Fig number | Pups*/experiment | | Dams | | Sex as variable | Assay kit Cat # | Supplier/ Reference |
|---|---|---|---|---|---|---|---|---|
| | | Con | Oxt | Con | Oxt | | | |
| DCE-MRI imaging | Fig 1A | | | 3 | 3 | No | | |
| R2* measurement | Fig 1B | | | 3 | 6 | No | | |
| Lactate colorimetric assay | Fig 2A | 8M/8F | 8M/8F | 8 | 8 | Yes | K-607 | Bio Vision |
| 4-hydroxynonenal assay | Fig 2B | 8M/8F | 8M/8F | 8 | 8 | Yes | STA-838 | Cell Bio Labs |
| Protein carbonyl assay | Fig 2C | 8M/8F | 8M/8F | 8 | 8 | Yes | STA-310 | Cell Bio Labs |
| Total glutathione assay | Fig 2D | 8M/8F | 8M/8F | 8 | 8 | Yes | STA-312 | Cell Bio Labs |
| RNA sequencing | Fig 3 | 5 | 5 | 5 | 5 | No | | |
| Behavioral study | Fig 4 | 12M/12F | 12M/12F | 9 | 9 | Yes | | |
| TaqMan qPCR array | Fig 5A | 8M/8F | 8M/8F | 8 | 8 | Yes | | |
| WB: ox-phos | Fig 5B | 5M/5F | 5M/5F | 5 | 5 | Yes | | |
| FcOIS imaging | Fig 6 | 7M | 5M | 7 | 5 | Yes | | |
| WB: HIF1a | Suppl | 3M/3F | 3M/3F | 3 | 3 | Yes | | |
| WB: CC3 | Suppl | 3M/3F | 3M/3F | 3 | 3 | Yes | | |

*One male and female pup/unique dam/treatment condition was used for experiments unless stated otherwise (behavioral study, fcOIS), with the dam as the experimental unit. M: male offspring; F: female offspring.

**Table 2:** TaqMan qPCR primers from ThermoFisher Scientific

| Gene | Taqman primers Cat# | Reporter dye | Formulation |
|---|---|---|---|
| 18S rRNA | Hs99999901_s1 | FAM-MGB | 20x |
| gapdh | Rn01775763_g1 | FAM-MGB | 20x |
| pgk1 | Rn01474008_gH | FAM-MGB | 20x |
| actb | Rn00667869_m1 | FAM-MGB | 20x |
| mt-cyb | Rn03296746_s1 | FAM-MGB | 20x |
| mt-nd1 | Rn03296764_s1 | FAM-MGB | 20x |
| mt-nd2 | Rn03296765_s1 | FAM-MGB | 20x |
| mt-nd4 | Rn03296781_s1 | FAM-MGB | 20x |
| mt-nd5 | Rn03296799_s1 | FAM-MGB | 20x |
| mt-atp6 | Rn03296710_s1 | FAM-MGB | 20x |
| mt-atp8 | Rn03296716_s1 | FAM-MGB | 20x |
| cat | Rn00560930_m1 | FAM-MGB | 20x |
| gpx1 | Rn00577994_g1 | FAM-MGB | 20x |
| gpx2 | Rn00822100_gH | FAM-MGB | 20x |
| gsr | Rn01482159_m1 | FAM-MGB | 20x |
| nox4 | Rn00585380_m1 | FAM-MGB | 20x |
| prdx1 | Rn00821587_g1 | FAM-MGB | 20x |
| sod1 | Rn00566938_m1 | FAM-MGB | 20x |
| sod2 | Rn00690588_g1 | FAM-MGB | 20x |
| srxn1 | Rn04337926_g1 | FAM-MGB | 20x |
| nqo1 | Rn00566528_m1 | FAM-MGB | 20x |
| bax | Rn01480161_g1 | FAM-MGB | 20x |
| bcl2l1 | Rn00437783_m1 | FAM-MGB | 20x |
| casp8 | Rn00574069_m1 | FAM-MGB | 20x |
| hif1a | Rn01472831_m1 | FAM-MGB | 20x |
| mt3 | Rn00588658_g1 | FAM-MGB | 20x |
| cygb | Rn00590627_m1 | FAM-MGB | 20x |
| nos2 | Rn00561646_m1 | FAM-MGB | 20x |
| mt-co1 | Rn03296721_s1 | FAM-MGB | 20x |
| mt-co3 | Rn03296820_s1 | FAM-MGB | 20x |
| nfkb1 | Rn01399572_m1 | FAM-MGB | 20x |

**Table 3:** List of antibodies and dilutions

| Antibody | Cat # | Supplier/ Reference | Stock concentration (mg/ml) | Working dilution |
|---|---|---|---|---|
| Mouse anti- Ox-phos ab | MS-604300 | Abcam | 1.5 | 1/1000 |
| Rabbit anti- HIF1a ab | 14179S | Cell Signaling Technology | 0.005 | 1/1000 |
| Rabbit anti- CC3 ab | 9661S | Cell Signaling Technology | 0.052 | 1/1000 |
| Rabbit anti- VDAC1 ab | ab-15895 | Abcam | 1 | 1/1000 |
| Mouse anti- Histone H3 ab | 9715S | Cell Signaling Technology | not determined | 1/1000 |
| Mouse anti- Beta actin ab | MA5-11869 | Thermo Fisher Scientific | 0.2 | 1/1000 |
| Goat anti- Rabbit IgG | 7074 | Cell Signaling Technology | not determined | 1/1000 |
| Horse anti- Mouse IgG | 7076 | Cell Signaling Technology | not determined | 1/1000 |

### 2. Supplemental Methods

**(i) Rapid sex determination of E21 offspring:** We developed a confirmatory test to establish the sex of E21 pups by performing a mini laparotomy to identify the presence (males) or absence (females) of testes. Testes can be identified as a globular structure covered with tortuous vasculature. Performed in conjunction with visual inspection of the anogenital distance, 100% accuracy can be readily achieved for sex identification.

**(ii) Behavioral experiments:**

**1 h open-field activity:** On P31 the OXT and CON rats were evaluated in terms of general ambulatory activity and exploratory behavior over a 1-h period in an open-field (41 x 41 x 38.5 cm high) constructed of Plexiglas and containing computerized photobeam instrumentation

(Kinder Scientific, LLC, Poway, CA). The apparatus included a frame that housed a 16 x 16 matrix of photocell pairs at ground level to quantify horizontal movement as well as another frame that contained 16 photocell pairs on its long axis, which was raised approximately 8.5 cm above the floor to measure vertical rearing behavior. General activity variables included total ambulations (whole body movements), vertical rearing frequency, and distance traveled in a 5.1 cm wide peripheral zone that surrounded the field. Classic measures of emotionality involved quantifying time spent in, distance travelled in, and number of entries made into a 10.2 x 10.2 cm central zone of the field.

**Social approach:** The day after being tested in the open field, the performance of the juvenile rats was assessed on the social approach measure, which was used to quantify sociability (tendency to initiate social contact with a novel conspecific), and preference for social novelty (tendency to initiate social contact with a novel versus a familiar conspecific). For the present studies we used our previously published protocol for mice, although the apparatus used was of a size scaled for rats. Specifically, the apparatus was a rectangular 3-chambered Plexiglas box (Stoelting Co., Wood Dale, IL.) with each chamber measuring 33 cm (w) x 100 cm (l) x 22 cm (h) containing Plexiglas dividing walls with square openings (10 x 10 cm). A withholding cylinder [30 cm (h); 16.5 cm diam] that contained vertical bars was used to sequester a stimulus rat. The presence of vertical bars allowed for minimal contact between rats but prevented fighting or sexual behaviors, and one was located in each outer chamber. A digital video camera connected to a PC computer that contained a tracking software program (ANY-maze, Stoelting Co., Wood Dale, IL.) that recorded the movement of a rat within the apparatus and quantified time spent, latency to enter, number of entries and distance traveled in each chamber, as well as time spent, latency to enter and number of entries made into an investigation zone surrounding each cylinder. The investigation zones were 20.5 cm in diameter, encompassing 2 cm around the withholding cylinders. An entry into the chambers was defined when a chamber contained 80% of the rat's body, while only the head was tracked in the zones surrounding the withholding cylinders to capture investigative social behaviors. Indirect lighting illuminated the test room and the entire apparatus was cleaned with Nolvasan solution while the cylinders were cleaned with 75% ethanol solution between tests.

Each test session consisted of 3 consecutive 10-min trials, the first trial being dedicated to habituation to the apparatus and general procedure, with the second two being test trials. For the first trial, each rat was placed in the middle chamber for acclimation to the entire apparatus. During the first trial, a rat was allowed to freely investigate and habituate to all three chambers, including the empty withholding cylinders. The second trial (test trial 1) involved placing an unfamiliar, sex-matched conspecific in one withholding cylinder, while the other was left empty, and the test rat was allowed to freely explore the apparatus and investigate the novel stimulus rat in the cylinder. For the third trial (test trial 2), the stimulus rat was left in the withholding cylinder in the same location while a novel rat was placed in the other cylinder and the test rat was allowed to explore the apparatus and investigate the two rats contained in the cylinders. Locations of the stimuli rats in the outer chambers for the test sequences were counterbalanced within and across groups. Entries, latency to enter, and time spent in each of the investigation zones and chambers were recorded for each rat.

**Elevated plus maze (EPM):** Three days after completing social approach testing, anxiety-like behaviors were evaluated in the EPM according to our previously described procedures for rodents. Given the young age and size of the juvenile rats in the study and the fact that we were using a 3-day protocol, the decision was made to use our mouse version of the apparatus instead of our apparatus that is used with adult rats. The entire apparatus was thoroughly cleaned with 70% EtOH prior to testing. Conditioned aversion to the open arms becomes strong across the 3 days of testing and we wanted to encourage at least a modest level of exploration of the open arms during the second and third test sessions to avoid "floor effects". The apparatus consisted of two opposing open arms and two opposing enclosed arms (36 x 6.1 x 15 cm) that extended from a central platform, (5.5 x 5.5 cm) which were constructed of black Plexiglas. The maze was equipped with pairs of photocells configured in a 16 (x-axis) x 16 (y-axis) matrix, the output of which was recorded by a computer and interface assembly (Kinder Scientific, Poway, CA, USA). A system software program (MotorMonitor, Kinder Scientific, Poway, CA, USA) enabled the beam-break data to be recorded and analyzed to quantify time spent, distance traveled, and entries made into the open and closed arms and center area. To adjust for differences in general activity, the percentage of distance traveled, time spent, and entries made in the open arms out of the totals (open arms + closed arms) for each variable were also computed. Test sessions were conducted in a dimly lit room with light being provided by two 13-watt black-light bulbs (Feit EcoBulbs). Each 5-min session began by placing a rat in the center of the maze and allowing it to freely explore the apparatus, with testing being conducted over 3 consecutive days.

**Observational fear learning (OFL):** On the two consecutive days following completion of EPM testing, empathy-like fear responses were assessed in the OFL task by conditioning the rats to context-dependent fear as a result of them observing a conspecific in brief distress (receiving foot shocks), using methods similar to previously published procedures (Keum et al., 2016). The apparatus (27cm x 31cm x 30.5cm) consisted of an acrylic chamber with a stainless-steel grid floor equipped with a house light, video camera, and was contained within a sound-attenuating chamber (Lafayette Instrument Co. and Actimetrics). The apparatus consisted of a custom clear acrylic stimulus chamber (13cm x 24.5cm x 20cm) which included a partition that separated the apparatus into two halves, and which contained 90, 1 cm diameter holes to allow for scent and sound penetration, and a grid floor was also present for delivery of shocks. A matte PVC cover was placed over the grid floor in the other half of the apparatus, designated the observer compartment, to prevent the test rat from receiving foot shocks. On day 1, a stimulus rat was placed in the stimulus chamber and an observer test rat was placed in the observer compartment. Baseline freezing behavior of the observer test rat was quantified for 5 minutes. During minutes 6-9, the stimulus rat received a 2 s 1.0 mA footshock every 10 s, and the freezing behavior of the observer test rat was quantified. On day 2, the observer test rat was again placed in the observer compartment and freezing behavior was quantified over the 4 min test to evaluate the contextual fear response in the absence of the stimulus rat. Freezing (no movement except for that associated with respiration) was quantified using FreezeFrame (Actimetrics, Evanston, IL), which allowed for the simultaneous visualization of behavior while adjusting a "freezing threshold" which categorized behavior as "freezing" or "not freezing" during 0.75 s intervals. The dependent variable analyzed from both trials was the percent of time spent freezing. Videos from each mouse were reviewed and some were eliminated from the statistical analyses because a rat may have been out of the field of view of the camera or the video may have become “locked” on a single frame making accurate measurement of freezing behavior not possible.

**Supplemental Figure 1**

Heatmap of significantly differentially expressed genes in OXT (top row) vs. saline (bottom row) exposed pups.

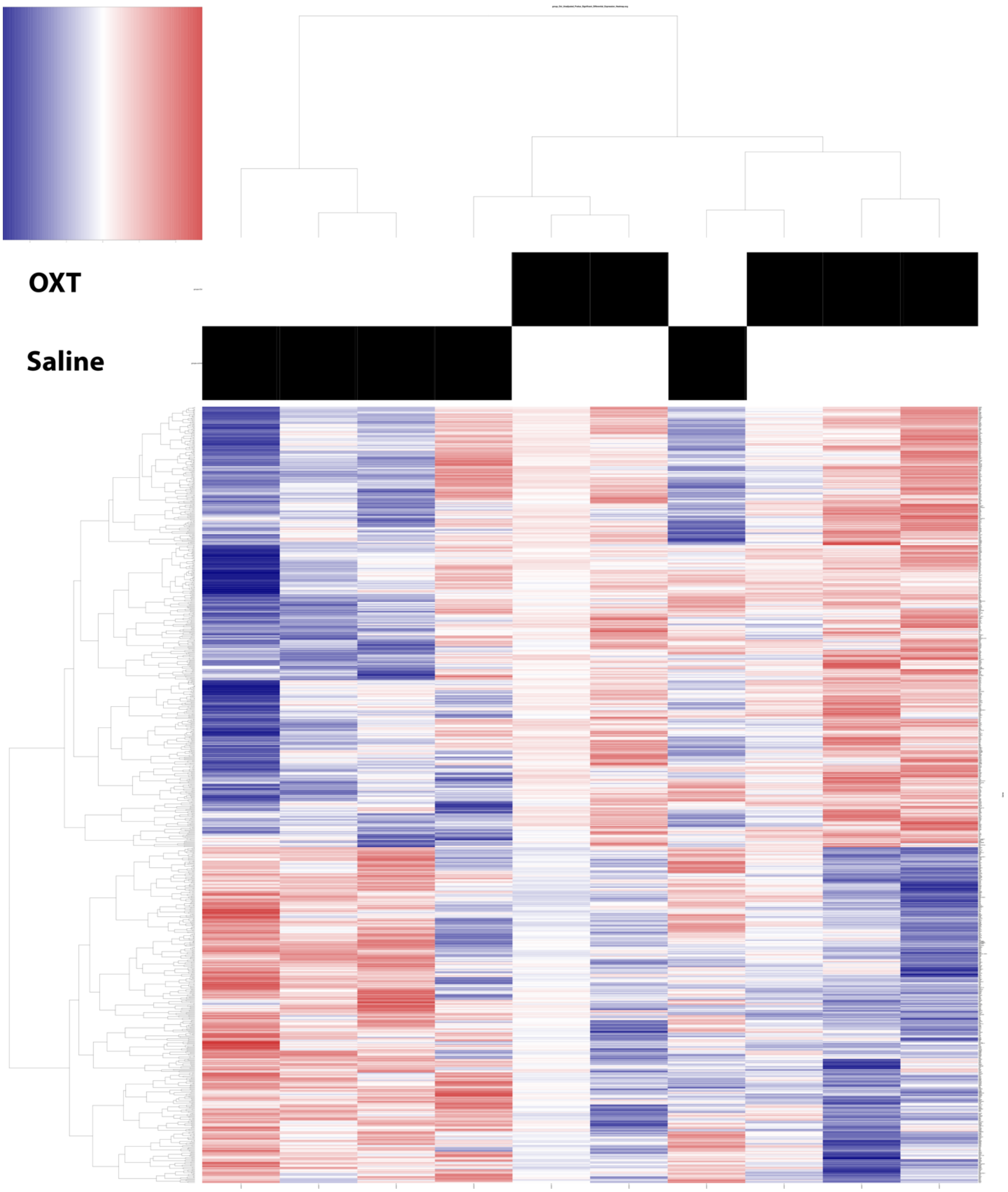

### Supplemental Figure 2

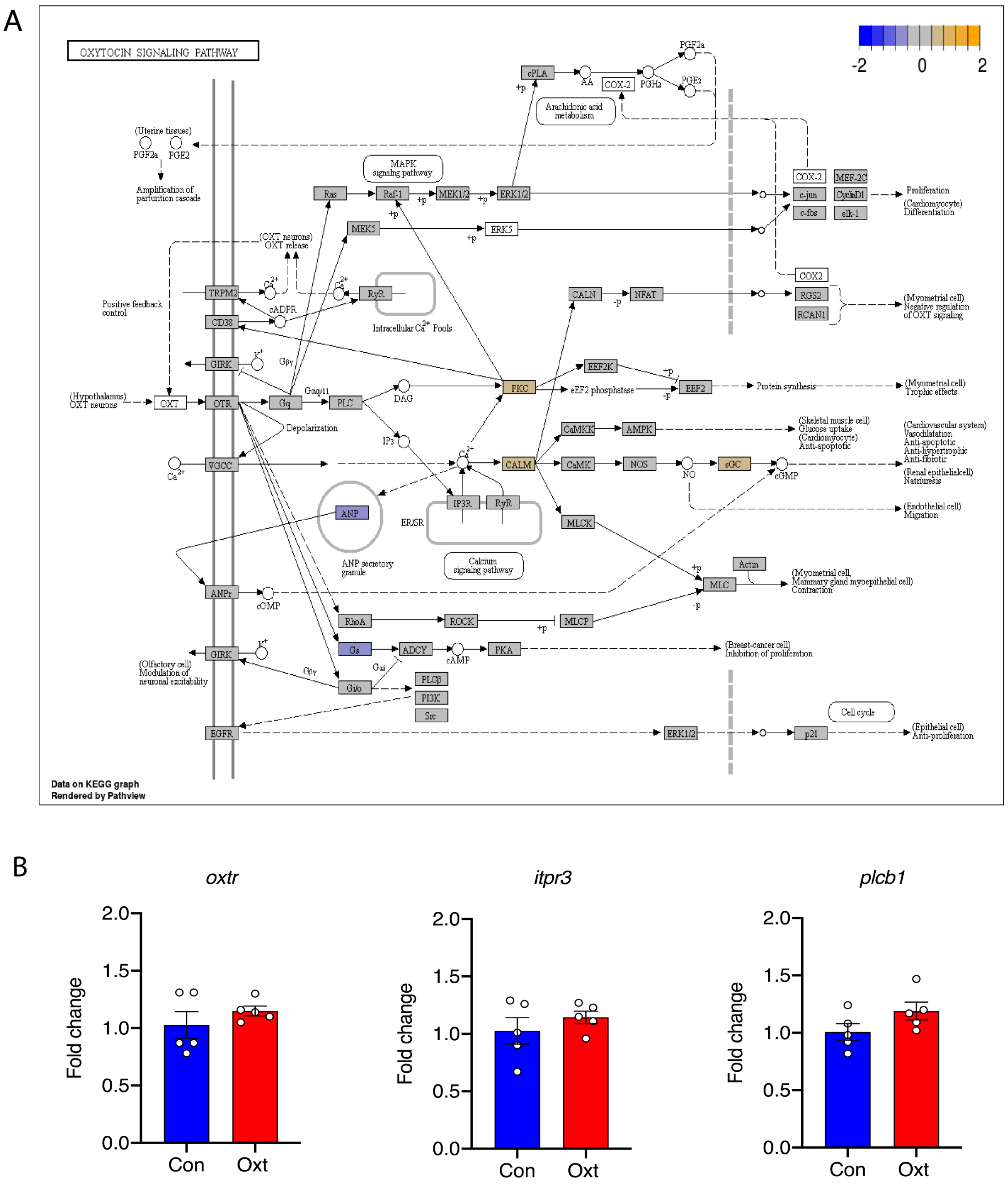


**Figure 2. Lack of significant involvement of the OXTR system.** Figure 2A shows the KEGG graph of the OXT signaling pathway in the fetal brain 24 h after OXT-induced aberrant uterine contractility. We pursued this with Taqman qPCR validation experiments for 3 genes integral to the OXT signaling pathway (*oxtr*, *itpr3*, and *plcb1*) and confirmed lack of significant differences between the control and OXT groups (Figure 2B) (n=5 per condition). Data were analyzed with student's t-test and presented as mean ± SEM.

**Supplemental Figure 3**

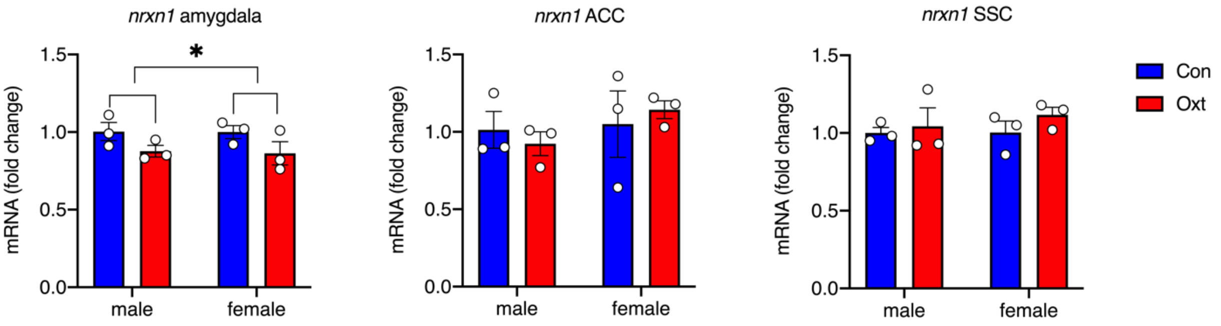


**Figure 3. *In utero* exposure to OXT-induced hypercontractility was associated with decreased *nrxn1* expression in the amygdala.** *Nrxn1* expression was assessed in the amygdala, anterior cingulate cortex (ACC), and the somatosensory cortex (SSC) of P28 male and female offspring exposed to either saline or OXT treatment *in utero* (one male and female pup per dam for n=3 dams per treatment condition). *Nrxn1* expression was reduced in the amygdala of both male and female offspring after OXT, but unchanged in the ACC and SSC. There were no sex x treatment interactions in any of the brain regions. Data were analyzed with 2-way ANOVA and presented as mean ± SEM; *p<0.05.

**Supplemental Figure 4**

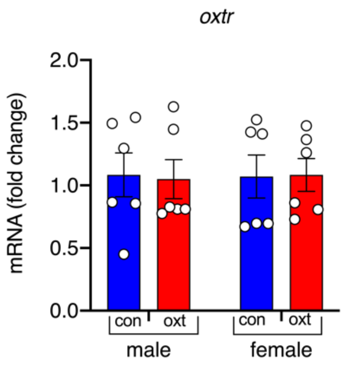


**Figure 4. OXT was not associated with sex-specific differences in fetal brain *oxtr* expression at 1 h.** To ensure that the changes observed in the fetal brain at E21 following OXT bolus in the dam was due to placental ischemia-hypoxia and not due to direct engagement with the oxytocin receptor (oxtr), we performed TaqMan® qPCR with a pre-validated *oxtr* probe (one male and female pup per dam per treatment condition for n=6 dams/treatment). We observed neither treatment nor sex differences in the expression of OXTR gene in the fetal cortex. This provided reassurance that the observed changes were not due to direct OXTR activation. Data were analyzed with 2-way ANOVA and presented as mean ± SEM.

**Supplemental Figure 5**

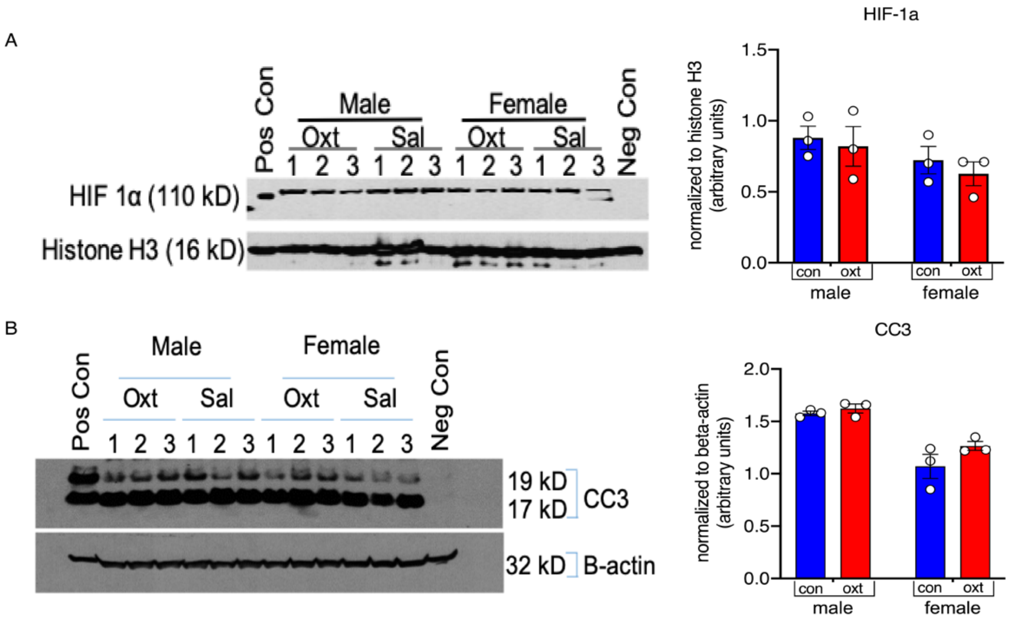


**Figure 5. 5A). Transient OXT-induced hypercontractility does not increase nuclear hif-1a protein expression in the developing fetal brain.** E21 fetal brains from both male and female pups were collected 1 h after OXT (100 mcg/kg) or saline (n=3 dams/treatment with one pup of either sex/dam). Hif1a protein expression in the nucleus, normalized to histone H3, was not different between groups. **5B). Transient OXT-induced hypercontractility does not induce apoptosis in the developing fetal brain.** E21 fetal brains from both male and female pups were collected 18 h after OXT (100 mcg/kg) or saline (n=3 dams/treatment with one pup of either sex per dam). Cleaved caspase-3 protein expression (19kD band), normalized to beta-actin, was not different between groups, though the overall level of cc3 expression was significantly lower in females of both treatment conditions. Data were analyzed with 2-way ANOVA and presented as mean ± SEM.

**Supplemental Figure 6**

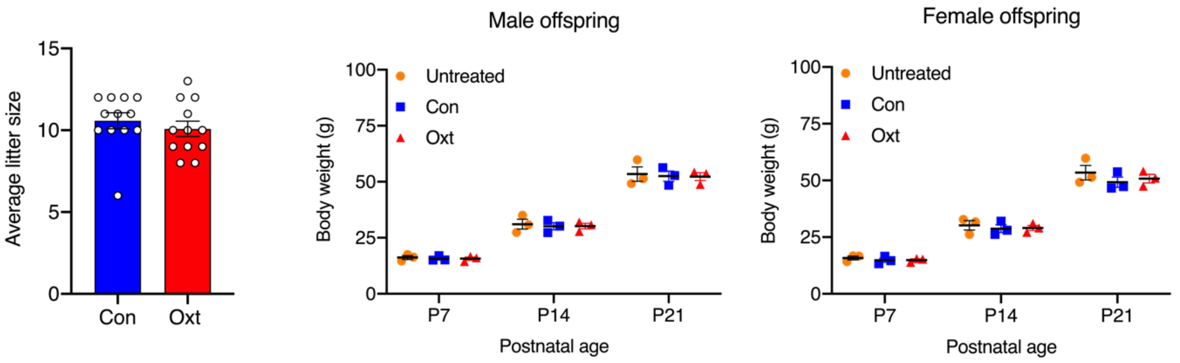


**Figure 6. Exposure to OXT-induced aberrant uterine contractility does not affect survival or growth of the offspring.** Average litter size was no different between the control and OXT-treated groups. At an OXT dose of 100 mcg/kg body weight, we did not observe any stillbirths. Litter size data were combined from behavioral and functional connectivity studies (n=12 dams per condition). In addition, we evaluated weight gain trajectories for male and female offspring for both control and OXT condition from data generated during the second cohort of behavioral studies (n=3 dams per condition). We chose this cohort because we measured body weights at 3 different time points (P7, P14, P21). Weight gain data were similar for both control and OXT-exposed male and female offspring, with sex-matched offspring from untreated dams included for comparison. Data were analyzed with 2-way ANOVA and presented as mean ± SEM.

**Supplemental Figure 7**

Group averaged correlation maps and matrices for saline and OXT-exposed male juvenile rats

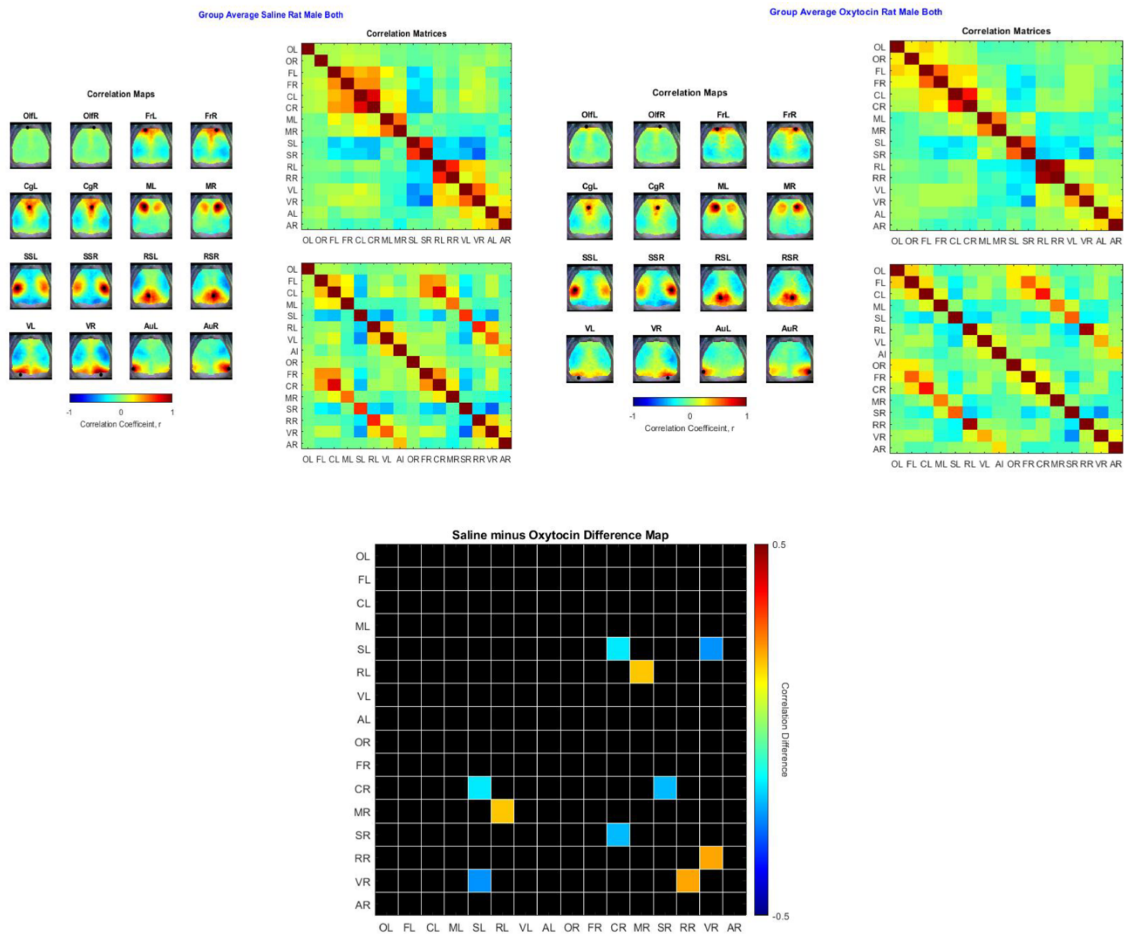


**Supplemental Figure 7.** Group averaged correlation maps for both saline (top left) and OXT-exposed (top right) offspring along with a difference map (below) showing regions where fc is variable. Abbreviations: L – left, R – right, Ol - olfactory, F – frontal, Cg – cingulate, M – motor, SS – somatosensory, V – visual, Au – auditory.
